# Development of an *ex vivo* model to study the zoonotic transmission of noroviruses

**DOI:** 10.1101/2025.10.28.685106

**Authors:** Nele Villabruna, Myra Hosmillo, Ian Goodfellow, Niklas Arnberg, Juan Bárcena, Silke Rautenschlein, Andreas Beineke, Gisa Gerold

**Affiliations:** Institute of Biochemistry and Research Centre for Emerging Infections and Zoonoses (RIZ), University of Veterinary Medicine Hannover, 30559 Hannover, Germany; Department of Pathology, University of Veterinary Medicine Hannover, 30559 Hannover, Germany; Division of Virology, Department of Pathology, University of Cambridge, Addenbrooke’s Hospital, Hills Road, Cambridge, CB2 0QQ, UK; Department of Clinical Microbiology, Umeå University, Umeå, Sweden; Umeå Centre for Microbial Research (UCMR), Umeå, Sweden; The Laboratory for Molecular Infection Medicine Sweden (MIMS), Science for Life Laboratory (SciLifeLab), Umeå University, Umeå, Sweden; Centro de Investigación en Sanidad Animal (CISA-INIA/CSIC), Madrid, Spain; Clinic for Poultry, University of Veterinary Medicine Hannover, 30559 Hannover, Germany; Institute of Virology, Medical University of Innsbruck, Innsbruck, Austria

## Abstract

Human noroviruses are the most common cause of viral gastroenteritis worldwide, causing sporadic cases and outbreaks. Genetically related animal noroviruses infect wild and domestic animals including pigs and dogs. To investigate the potential of noroviruses to jump the species barrier, we used porcine, canine, and avian precision-cut intestinal slices. We show that human norovirus can bind to pig and dog intestinal tissue and be taken up with a similar efficiency as the porcine and canine noroviruses, respectively. In contrast, no binding or internalization of human, porcine, or canine noroviruses was detected in chicken tissue. We further showed that human norovirus can replicate to some degree in pig intestinal tissue, infecting single cells. In contrast, while animal noroviruses attached to human intestinal cells, intracellular uptake was limited. This suggests that human-to-animal transmission is more likely than animal-to-human transmission and that viral uptake likely presents a species barrier.

## Introduction

*Norovirus* is a genus of the *Caliciviridae* family and the most common non-bacterial cause of foodborne gastroenteritis worldwide. The human norovirus single-stranded positive RNA genome is composed of three open reading frames (ORF1-3) of which ORF2 is encoding the major structural protein which makes up the viral capsid [1]. Based on capsid amino acid similarity, noroviruses are categorized into ten genogroups (GI–GX) that are further divided into 48 genotypes [2]. Viruses within GI, GII, GIV, GVIII and GIX are commonly found in humans, with GII.4 variants having caused at least six pandemics in the last 20 years. GII.4 variants also cause the majority of outbreaks and sporadic infections [3, 4]. Viruses of the other genogroups have been detected in a variety of wild and domestic species and are considered species-specific. These include cattle, sheep, and yaks (GIII), cats and dogs (GIV, GVI and GVII), rodents (GV), bats (GX) and some unclassified genogroups were found in harbour porpoises and Californian sea lions [2]. Exceptions are the porcine genotypes (GII.11, GII.18, and GII.19) and the canine genotype GIV.2, which cluster in the otherwise human norovirus genogroups GII and GIV, respectively. This close genetic relatedness between animal and human noroviruses raises the question whether these noroviruses could potentially be transmitted between humans and animals.

The detection of human sera IgG that recognize canine norovirus virus-like particles (VLPs) suggests that there is some exposure to animal noroviruses [5, 6]. However, no animal norovirus has been reported in symptomatic or asymptomatic individuals to date. In contrast, more studies have reported antibodies recognizing human norovirus VLPs in mammals, primarily in pigs and dogs, but also some primates. These findings could not only be explained by cross reactivity with animal noroviruses and likely stem from exposure to human norovirus (reviewed in [7]). In addition, several studies have reported the presence of GI and GII human norovirus RNA in stool samples of wild and domestic animals, most commonly in pigs [8]. These sequences shared a 94 - 100% nucleotide identity with noroviruses detected in humans. Especially with GII.4 variants the detection in animals postdated the dominant circulation of these variants in the human population indicating human-to-animal spillover events. Since these studies usually do not include human samples, only two studies showed a direct link that included samples of dogs and their owners from the same time period, demonstrating the same human norovirus in humans and animals [9, 10]. Caliciviruses infect a range of non-mammalian species, including reptiles, fishes, amphibians, and avian species (e.g. chickens) but noroviruses seem restricted to mammalian species [11]. Nevertheless, two studies have reported human norovirus RNA in bird droppings [12, 13].

Regarding potential barriers of cross-species transmission, human experimental infections and *in vitro* studies have shown that human noroviruses use histo-blood group antigens (HBGAs) as attachment factors [14–19]. HBGAs are expressed on intestinal epithelium and are also secreted in bodily fluids like saliva and milk [20–22]. A subset of the human population, so called non-secretors, do not express HBGAs in their intestine and are resistant to infection with some noroviruses like GII.4 [16–18, 23]. HBGAs are likely broadly expressed among mammalian species [24, 25] and have been described to also be attachment factors for canine and bat noroviruses [26, 27]. While HBGAs are necessary for norovirus attachment they are not sufficient for infection. For example, norovirus can attach to HEK293T and Huh-7 cells overexpressing HBGAs and when these cells are transfected with viral RNA, they support genome replication, however no full infectious cycle occurs [28–30]. This indicates that noroviruses readily bind to tissue, but host restriction might occur downstream.

To investigate norovirus attachment, uptake and replication across species, we investigated the host barrier in porcine, canine and avian hosts. Pigs and dogs were selected for this study because most evidence points to them as potential reservoirs for human noroviruses. Moreover, they are common farm animals and pets that live in close proximity to people, providing ample opportunity for spillover. Chickens were included as control species, as there is little evidence of human or animal norovirus infection in poultry. To study the potential of human noroviruses to infect various animal species *in vitro* we used precision-cut intestinal slices (PCIS). Although precision-cut tissue slices are more commonly used in toxicological and pharmaceutical research, they have also been used to study pathogen host interactions [31–36].

## Results

### Setting up precision cut-intestinal slices of pig, dog and chicken origin

To establish PCIS from three animal species, fresh intestinal tissue of pig, dog and chicken origin was obtained and cut as described below. PCIS viability was tested at 0 h and 24 hours post preparation by measuring ATP levels over tissue mass. PCIS viability varied within, but also between species in all three species. The ATP levels in pig and dog PCIS dropped after 24 h to half of the starting concentration, but in chickens it increased to approximately double the starting levels (**Figure 1A**). Despite this drop, the levels were above the 2 pmol ATP/µg protein threshold for tissue viability, as previously defined [37, 38]. The macrostructure of the tissue 24 h post cutting consistently showed intact villi and crypts by haematoxylin & eosin and immunofluorescent staining (**Figure 1B**). These looked comparable between species but are exemplarily shown here for pigs.

**Figure 1.**
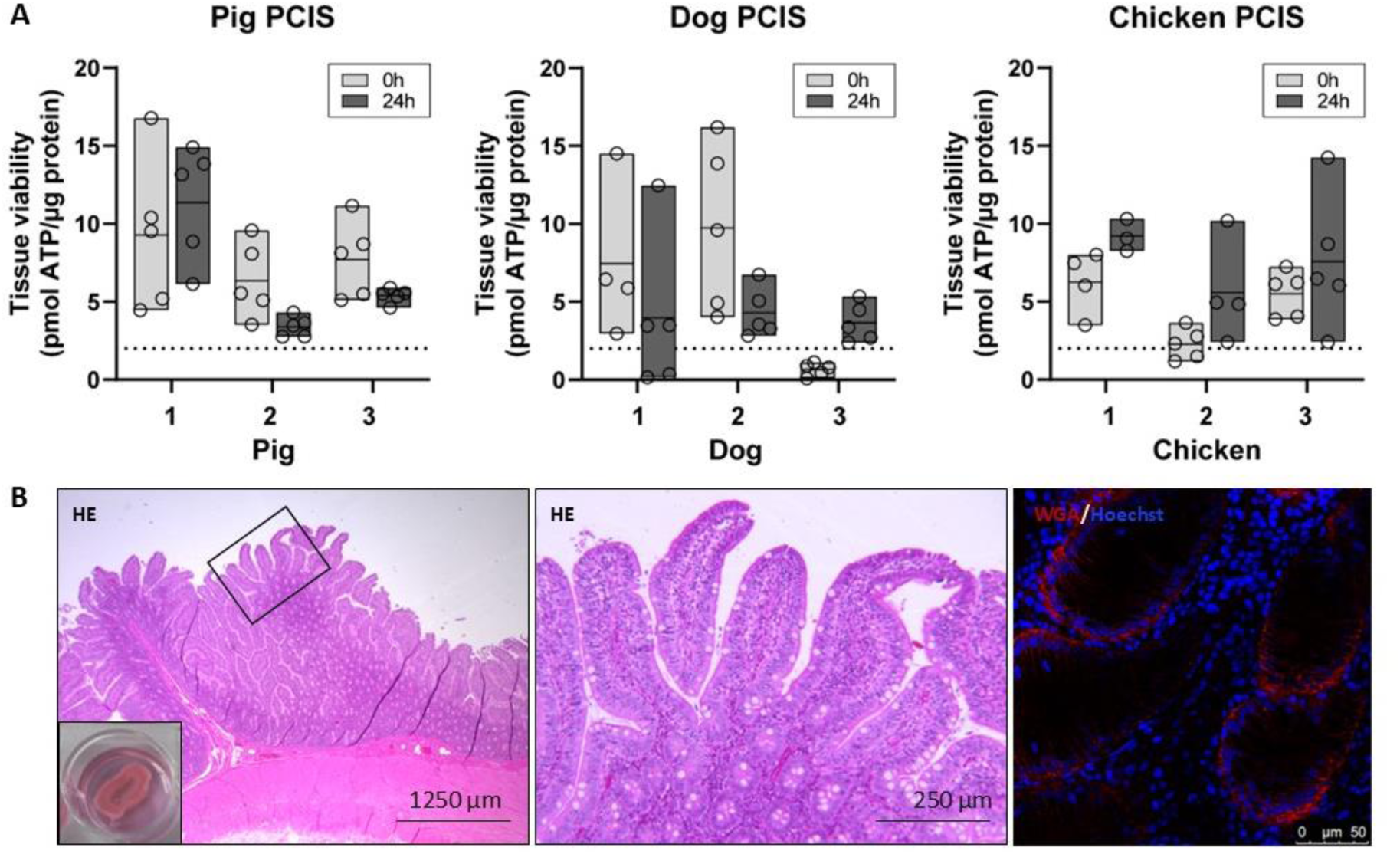
Precision-cut intestinal slices (PCIS) from fresh pig, dog, and chicken small intestinal tissue remain viable for up to 24 hours. (**A**) PCIS were prepared by embedding fresh dog, pig, and chicken intestinal tissue in low-melting agarose. The embedded tissues were cut into 250-µm slices and kept in culture medium for 24 h. Tissue viability was measured by quantifying the ATP concentration relative to tissue mass (total protein content (µg)). The dotted line indicates the cut-off of 2 pmol ATP/μg protein [37]. At least four PCIS were tested per timepoint and animal. Data of three individuals are shown per species. (**B**) Haematoxylin & eosin (HE) and immunofluorescent staining of pig PCIS collected from a jejunal section of the small intestines was performed to visualize tissue integrity 24 hours after cutting. The inset depicts a PCIS in culture medium. Nuclei were stained with Hoechst 33342 (blue) and cell membranes with wheat germ agglutinin (WGA, red).

### Human norovirus VLPs bind to and are internalized into tissue of pig and dog origin

Human norovirus (HuNoV) attaches to the epithelium of formalin-fixed, paraffin-embedded (FFPE) intestinal tissue of a broad range of species [25]. To ensure that the epithelium was preserved after cutting and could still be recognized by norovirus, the binding ability and subsequent internalization of human GII.4 Dijon, porcine GII.11, and canine GVI.2 norovirus (**Figure 2**) were tested in pig, dog, and chicken intestinal tissues using FFPE PCIS. To that end, we generated virus-like particles (VLP), i.e. particles without encapsidated genome and labelled them with a fluorophore (AF488). Additionally, since several norovirus genotypes, including HuNoV GII.4, are HBGA-dependent for infection of human organoids [19], samples were phenotyped according to their secretor status (α1,2 fucose group) using *Ulex europaeus* agglutinin 1 (UEA1) lectin staining.

**Figure 2.**
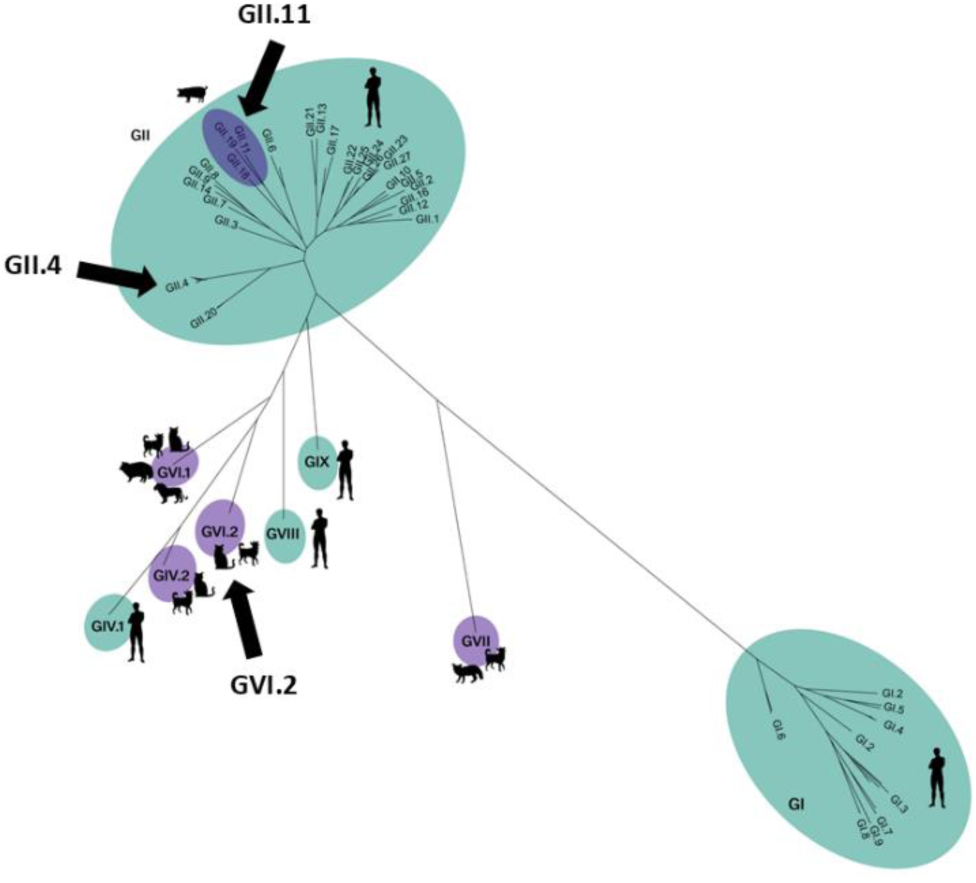
Maximum likelihood tree based on the whole VP1 sequence. Depicted are norovirus genogroups and genotypes infecting humans, pigs, and carnivores. Porcine and canine/feline noroviruses group within the same genogroups as human noroviruses, GII and GIV, respectively. The genotypes used for VLP binding in this study are marked with arrows.

Bound AF488-labelled VLPs to PCIS of pig, dog, and chicken intestine were visualized by the avidin– biotin complex method. HuNoV GII.4 VLPs attached to the epithelium of all pig and dog PCIS and additionally to goblet cells in pig but not to chicken PCIS (**Figure 3**). As expected, canine VLPs attached to 7/7 dog PCIS, while porcine VLPs attached to 2/7 pig PCIS. For the latter, the signal was not as strong as with GII.4. Attachment of HuNoV VLPs and canine VLPs were distributed along villi and crypts, while porcine VLPs were only observed in the villi. Neither the human nor animal norovirus VLPs attached to any of the chicken tissues. Furthermore, pig and dog tissues, but not chicken, tested positive for α1,2 fucose indicating that these samples are secretor positive.

**Figure 3.**
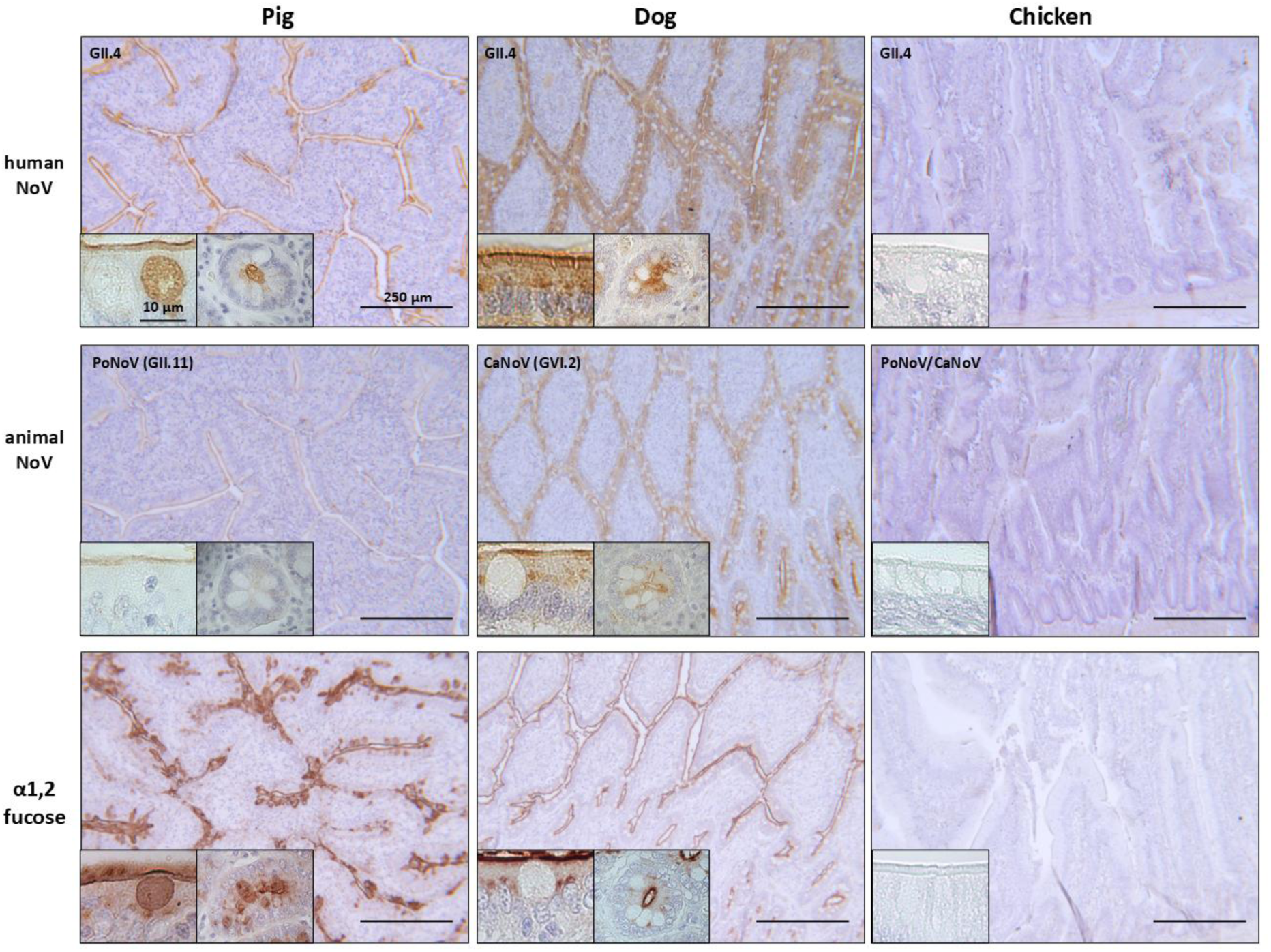
HuNoV VLPs attach to porcine and canine PCIS, but not of galline PCIS. Alexa Fluor 488-labeled norovirus VLPs were tested for their ability to bind to formalin-fixed, paraffin-embedded PCIS. As binding controls, porcine (PoNoV, GII.11) and canine (CaNoV, GVI.2) norovirus VLPs were tested on PCIS of their respective hosts. The secretor status of the tissues was confirmed by *Ulex europaeus* agglutinin 1 (UEA1) lectin staining. Brown staining depicts attached VLPs or the presence of α1,2 fucose group by lectin staining, nuclei are stained in blue (haematoxylin). Seven individual PCIS were tested per species. Images shown are representative. Scale bars: 250 µm, insets 10 µm.

In addition, VLP attachment and internalization were corroborated in live tissue using the inside-out assay that allows to differentiate between attached and internalized VLPs (**Figure 4A**) [39]. Human GII.4 VLPs attached to and internalized into PCIS of pig and dog origin but not of chicken origin. (**Figure 4B-G**). Porcine GII.11 and canine GVI.2 VLPs also attached to and were internalized into pig and dog PCIS, respectively. In pigs, 65 – 85% of HuNoV VLPs was internalized compared to 83 – 100% of porcine VLPs (**Figure 4C**). In dogs, 89 – 93% of HuNoV VLPs was internalized compared to 65 – 71% of canine VLPs (**Figure 4E**). When heating VLPs to 95°C for 10 min, binding was eliminated.

**Figure 4.**
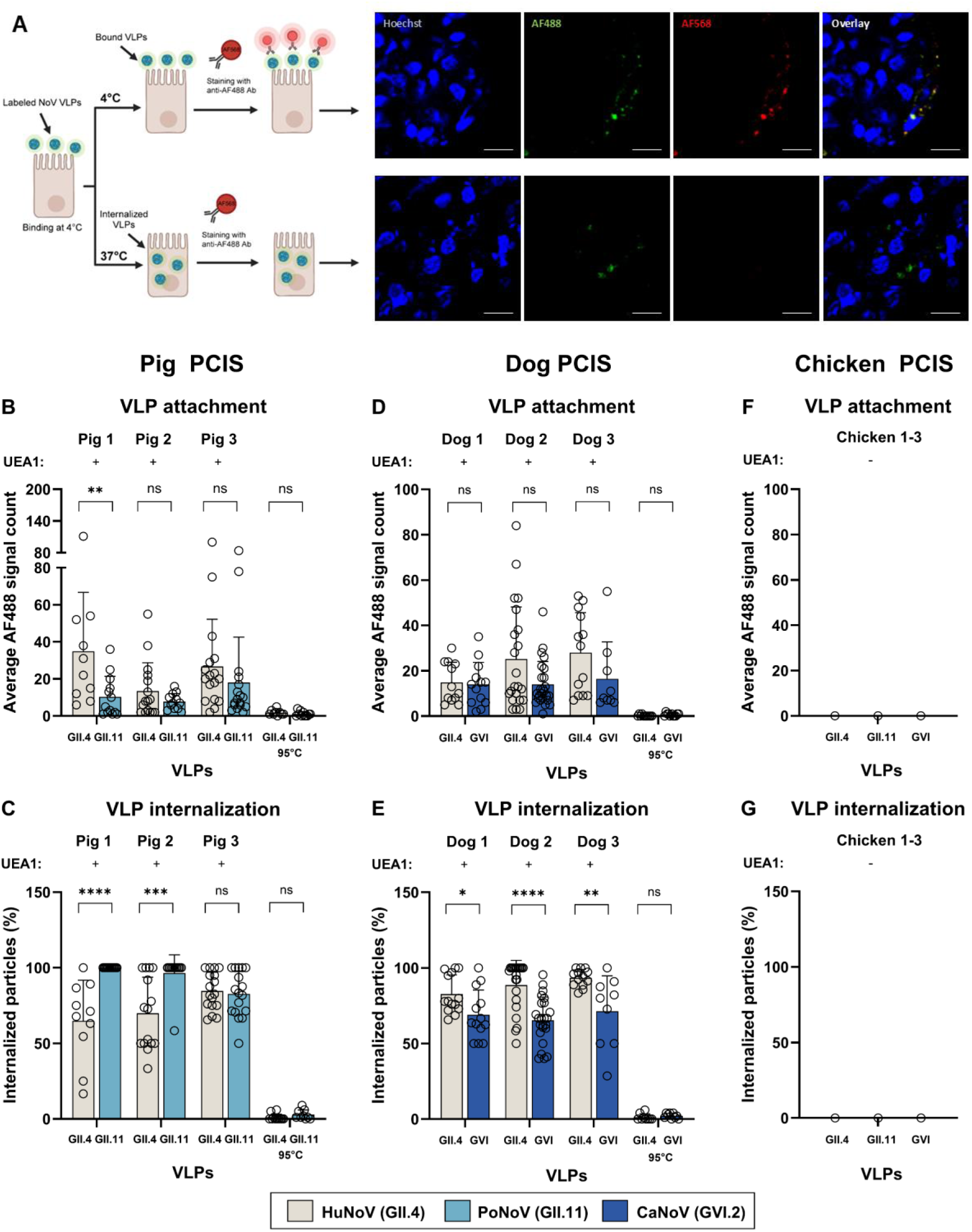
Human norovirus is efficiently taken up into PCIS of pig and dog but not chicken origin. HuNoV VLP binding and internalization was tested in an inside-out assay using live PCIS. (**A, left image**) Modified schematic diagram of inside-out assay was created in BioRender as described elsewhere [39]. (**A, right image**) AF488-labeled norovirus VLPs (green) were allowed to bind to animal PCIS on ice (4°C), preventing VLP internalization. An anti-AF488 antibody (AF568-labeled) was added to mark extracellular VLPs, which appear red in addition to the green label. When the temperature was increased to 37°C, VLPs were internalized and no longer accessible to antibodies. Therefore, a single green signal indicates internalized VLPs, while a double green and red signal visualizes extracellular, cell surface-bound VLPs. Nuclei were stained with Hoechst 33342. Representative immunofluorescent images are shown. (**B, D,** and **F**) To quantify VLP attachment, AF488 signal per image of the binding control was counted. (**C, E,** and **G**) To quantify VLP internalization, the percentage of internalized VLPs (only AF488-positive) versus all VLPs (all AF488-positive events) was calculated. The secretor status of the tissues was confirmed by *Ulex europaeus* agglutinin 1 (UEA1) lectin staining, as indicated on the top of the graph. As control, VLPs were heat inactivated for 10 min at 95°C. Per species, three individuals were tested. Mann-Whitney test was used to test for significance: **** = P ≤ 0.0001, *** = ≤ 0.001, ** = P ≤ 0.1, * = P ≤ 0.05, ns = P > 0.05. Scale bar = 10 µm in panel A.

### Animal noroviruses attach to but are not internalized into human intestinal cells

Some serological evidence suggests that humans could be exposed to animal noroviruses [5, 6]. We, therefore, investigated the ability of animal noroviruses to attach to and internalize into human intestinal cells. As no human biopsies were available to produce PCIS, the human duodenal cell line HuTu80 was used. Surprisingly, porcine (GII.11) and canine (GVI.2) norovirus VLPs attached significantly better to HuTu80 cells with 74 and 50 average AF488 signal counts, respectively compared to HuNoV VLPs (GII.4 Dijon) with 13. However, only 17% porcine VLPs and 16% canine VLPs were internalized, compared to 87% uptake of human VLPs (**Figure 5A** and **B**).

**Figure 5.**
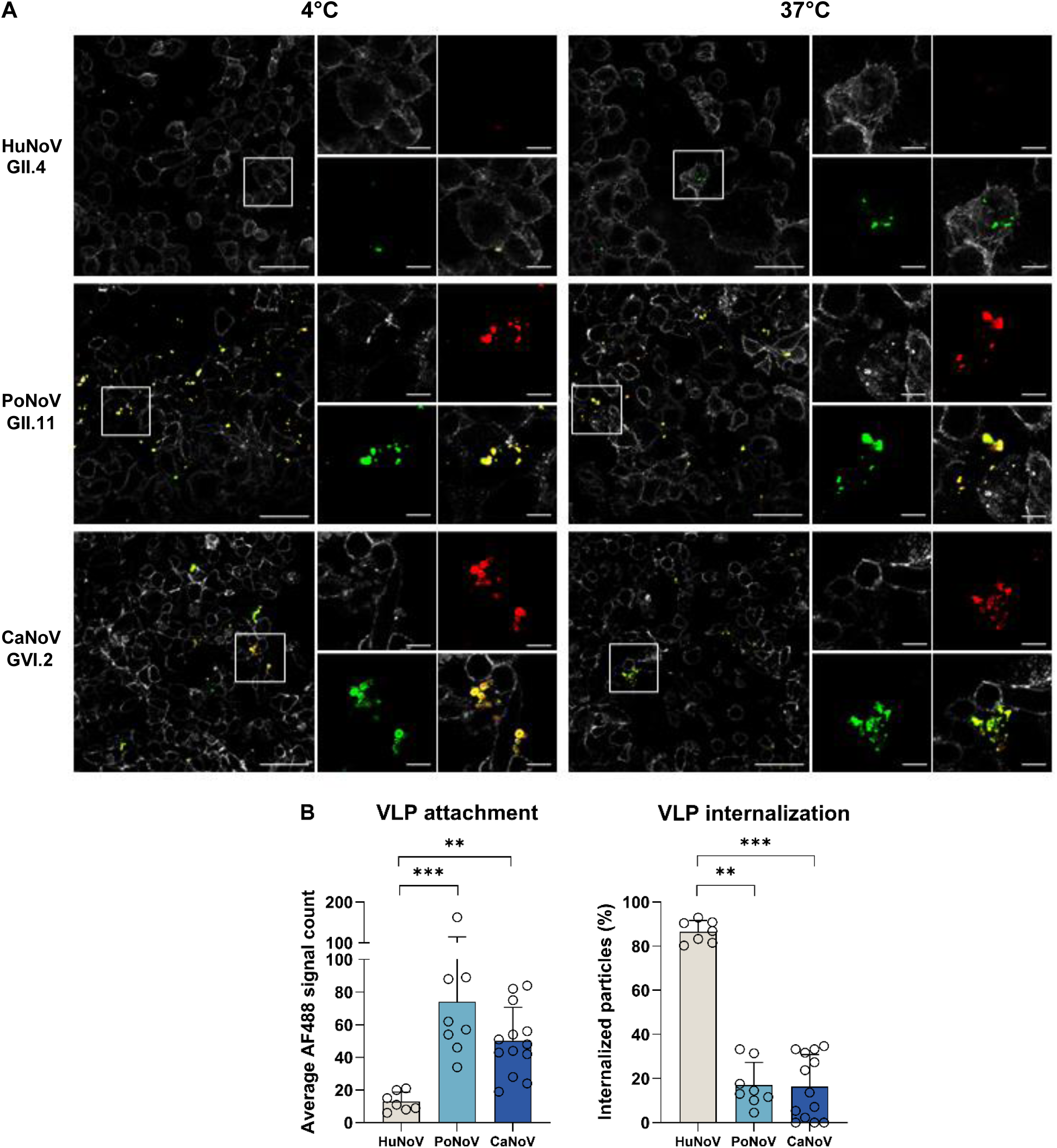
Animal and human norovirus attachment and internalization into human intestinal cells. Human and animal norovirus VLPs were allowed to attach and internalize into HuTu80 cells using the same assay as described in Figure 4. (**A**) Immunofluorescent (IF) images of AF488-labelled human (HuNoV, GII.4), porcine (PoNoV, GII.11), and canine (CaNoV, GVI.2) VLPs at 4°C (binding control) and 37°C (internalization). Membranes are stained with AF647-conjugated wheat germ agglutinin (**B**) Quantification of IF images. Significant differences between VLP attachment and internalization of PoNoV and CaNoV and that of HuNoV was tested using One-way Anova, Kruskal-Wallis Test. **** = P ≤ 0.0001, *** = ≤ 0.001, ** = P ≤ 0.1, * = P ≤ 0.05, ns = P > 0.05. Scale bar = 50 µm, insets = 10 µm.

### Human norovirus replicates in pig PCIS

To examine whether internalization of HuNoV live virus into porcine PCIS can lead to norovirus replication, GII.4 Sydney[P16] live virus was used, alongside transmissible gastroenteritis virus (TGEV), a porcine intestinal coronavirus as a relevant control virus. After 24 hours we detected an increase in viral genome copy numbers for HuNoV GII.4 (**Figure 6A**) and porcine TGEV (**Figure 6B**) with the highest increase of 143-fold and 344-fold, respectively. By IFA, using antibodies against dsRNA and viral capsid (HuNoV), or nucleoprotein (TGEV), positive cytoplasmic foci can be detected in individual cells, suggesting the presence of HuNoV and TGEV replication in pig PCIS. There was no signal at timepoint 0h and in mock-infected tissues.

**Figure 6.**
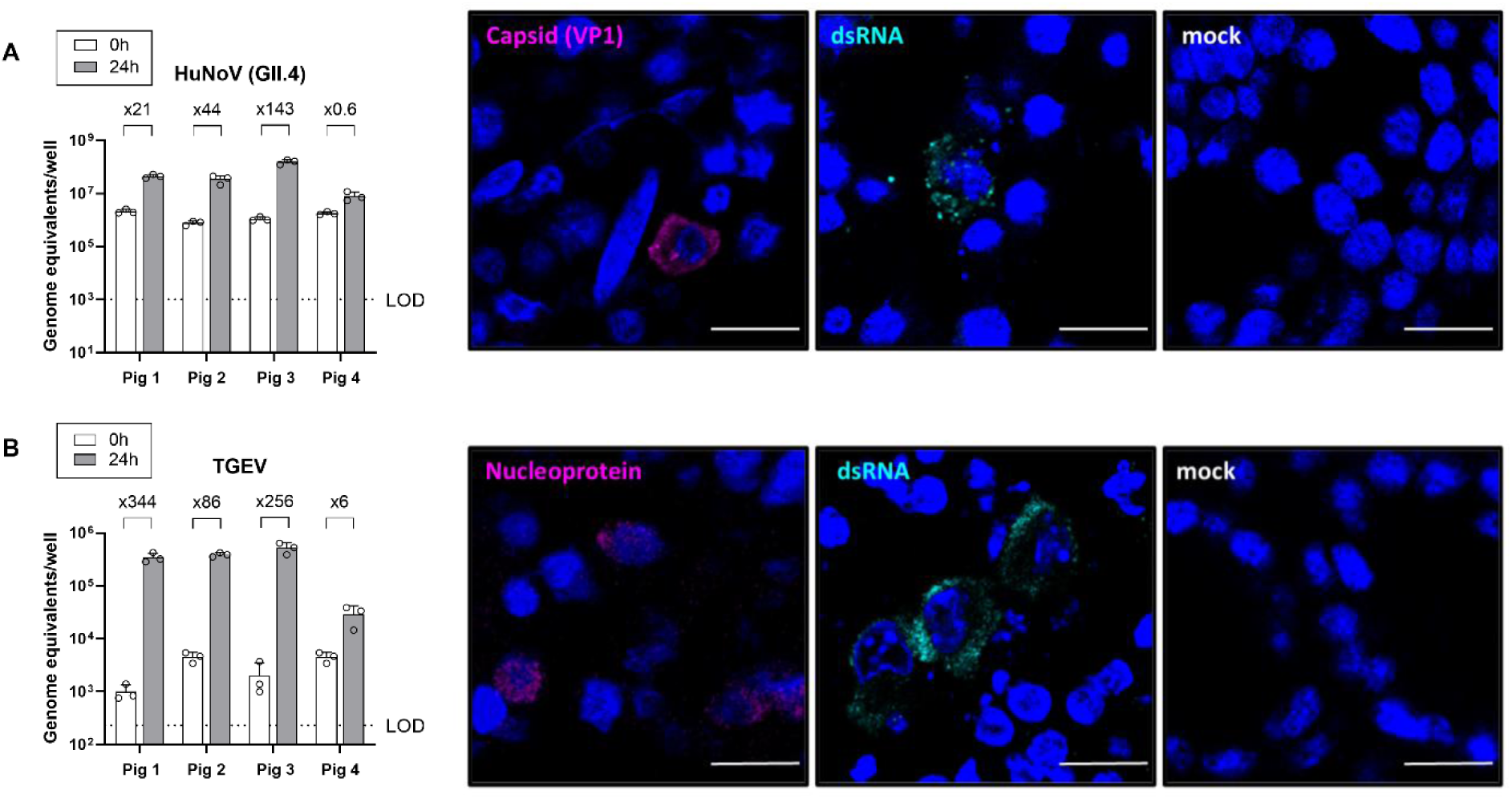
Human norovirus and transmissible gastroenteritis virus (TGEV) replicate in pig PCIS. Pig PCIS were inoculated with (**A**) HuNoV GII.4 Sydney[P16] or (**B**) TGEV PUR46 strain and incubated for 1 h at 37°C. Unbound virus was removed, PCIS were washed, fresh medium was added and PCIS were incubated for 24 h at 37°C. The supernatant was harvested right after the washing step for time point 0 h and after 24 h. GII.4 and TGEV RNA was detected by RT-qPCR with TGEV- or HuNoV-specific primers and probes. LOD = limit of detection. For immunofluorescence, HuNoV-inoculated pig PCIS were stained with antibodies recognizing the major capsid protein (VP1, purple) and dsRNA (cyan). TGEV-inoculated pig PCIS were stained with antibodies against the nucleoprotein (purple) and dsRNA (cyan). Nuclei were stained with Hoechst 33342 (blue). Scale bar = 10 µm.

## Discussion

Noroviruses infect a broad host range including domestic animals like pigs, dogs, and cows and wild animals like bats, harbour porpoises, and rodents, but the possibility of inter-species transmission remained largely elusive [2]. Here we set out to investigate whether noroviruses can jump the species barrier. We chose pigs and dogs as previous literature points to them as potential reservoir for human norovirus infection. Chickens served as a control species as there is no link between noroviruses and poultry. GII.4 attachment to pig and dog PCIS was comparable to that of host specific PoNoV and CaNoV, respectively. In contrast, none of the VLPs attached to the chicken tissues. These results matched the secretor status as all pig and dog but none of the chicken tissues were secretor positive. Our findings support results of previous studies using FFPE tissues where GII.4 attached to a broad range of animal species but not chickens or other avian species and this correlated with the presence of the α1,2 fucose group on the intestinal epithelium [25, 27].

Although HBGAs play an important role in host susceptibility, norovirus interaction with HBGAs alone does not necessarily lead to internalization and virus replication. For example, HBGA-like carbohydrates are also found on bacteria that naturally occur in human stool and norovirus attaches to these non-eukaryotic cells without infecting them [40, 41]. Similarly, human norovirus attaches to plant cells, especially those from lettuce [42, 43]. Attachment presumably occurs via HBGA-like carbohydrates, but it is very unlikely that norovirus particles are taken up, as they were detected exclusively on the outer bacterial membrane and cell wall. With regard to foodborne infections, oysters play an important role in norovirus outbreaks. They express HBGAs similar to humans thereby bioaccumulating norovirus particles but are not able to sustain infection [44, 45]. Similarly, the human intestinal cell line Caco-2 expresses HBGAs, but efforts to reliably infect these cells have not yielded consistent results [46, 47].

When we tested for internalization of human norovirus into animal tissue, we found high internalization of >65% in pig and dog tissue, which was comparable to that of the respective animal noroviruses. In a study by White et al. norovirus VLPs were tested for their ability to bind to 10 different mammalian and non-mammalian cells lines [46]. And while they bound to all included cell lines, most efficient uptake was seen into differentiated Caco-2 cells, into which 19% of VLPs were internalized. In comparison, only 2% were taken up into insect cells. The efficient VLP uptake that we detected suggests that pig and dog intestinal tissue can readily take up norovirus and that PCIS are a better model to study norovirus entry than Caco-2 cells. It should however be noted that we used a different genotype in this study, which could also explain a higher uptake rate. We used VLPs of an GII.4 Dijon strain which differs from the Sydney 2012 variant that we used for infection experiments. However, we think that the binding and internalization results are not variant specific, as GII.4 variants share their ability to recognize HBGAs and infect organoids [48, 49]. In contrast to the comparable uptake rate of human and animal norovirus into animal tissues, we saw a higher discrepancy in human intestinal cells; although surprisingly porcine and canine norovirus attached significantly better to human cells than GII.4, their uptake was inefficient compared to that of human norovirus (17% vs 87%). This suggests that human noroviruses are more likely to be transmitted to animals than vice versa, which is supported by the lack of cases detecting porcine or canine norovirus infection in humans. It was out of the scope of this study to further investigate through which pathway noroviruses are endocytosed into animal tissue. However, a previous study in organoids has shown that noroviruses enter via an endosomal acidification-dependent clathrin-independent carriers (CLIC) pathway [50].

Finally, we could show that human norovirus could replicate to some degree in pig PCIS, infecting single cells and the nature of these cells will be further investigated. As expected, replication was less efficient than what is reported form the organoid model [49, 51]. Not all tissues supported replication, and we do not know yet where this variation stems from. Since all animals were secretor positive it is likely an HBGA-independent factor. To our knowledge this is the first study showing that human norovirus can replicate in non-human tissue.

In conclusion, HBGA are a crucial factor to initiate norovirus attachment, potentially explaining the lack of binding to chicken intestine. While HBGA expression is necessary for norovirus attachment, it is not sufficient to render a cell line susceptible or even permissive to norovirus infection as some cell lines that do express HBGA do not support norovirus replication [47, 52]. It is likely that HBGA-dependent norovirus attachment does not necessarily initiate virus uptake indicating the presence of additional susceptibility factors. The nature of co-factor(s) is currently unknown, but an additional protein receptor has been hypothesized to play a role. The high uptake of human norovirus into animal PCIS and the relatively low uptake of animal norovirus into human cells suggests that host restriction occurs between virus attachment and internalization. Hence, norovirus entry should be further investigated. Taken together norovirus transmission from human-to-animal seems more likely than animal-to-human. Whether human noroviruses can be transmitted within an animal reservoir and be eventually be transmitted back to humans, should be further investigated.

## Material and Methods

### Ethics statement

A study approval from an ethics committee was not required, as no additional animal experiments were conducted.

### Cells and viruses

HuTu80 cells were cultured in Dulbecco’s modified Eagle medium (DMEM, Sigma Aldrich, St. Louis, MO, US), supplemented with 100 U/mL penicillin and 100 µg/mL streptomycin (Gibco, Waltham, MA, US) and 20 mM HEPES (Thermo Fisher Scientific, Waltham, MA, US) and 10% fetal bovine serum (Capricorn Scientific, Ebsdorfergrund, Germany) at 37°C and 5% CO_2_.

Stool samples from norovirus-positive individuals were received from the Robert Koch Institute and all typed as GII.4 Sydney]P16]. A 10% stool solution was prepared in PBS. The samples were sequentially centrifuged at 4°C for 10 min: at 500 x g to 5000 x g in 1000 g increments. TGEV was kindly provided by Paul Becher and Luis Enjuanes.

### Intestinal tissues

For dogs, intestinal tissue was removed during routine necropsy service at the Department of Pathology (1 to 4 h postmortem). Dogs with intestinal pathology were excluded. Dogs varied in ages, sex and breed. No animals were killed for the purpose of this study and written consent was given by the owners. For pigs, intestinal tissues were received from healthy six months-old animals from an abattoir. Chicken tissue was received from specific-pathogen-free healthy animals (Valo, BioMedia GmbH, Osterholz-Scharmbeck, Germany) used for other experiments (females, aged 6-7 month, under the permit TiHo-T-2022-16). Of each animal, stool sample was isolated from intestine and stored at -70°C. Those used for infection experiments were tested for presence of noroviruses by RT-qPCR.

### Preparation and culturing of precision-cut intestinal slices (PCIS)

PCIS preparation was based on the protocol by de Graaf et al. [37]. Fresh intestinal tissues were harvested from pigs, dogs, or chickens and put into precooled ROTI®Cell Hanks’ BSS with Magnesium and Calcium (Carl ROTH, Karlsruhe, Germany) on ice. Tissues were cut into 4-cm pieces and flushed several times with cold KHB buffer (NaCl 7.00 g/L (Carl ROTH) KCl 0.37 g/L (Carl ROTH), KH_2_PO_4_ 0.16 g/L (Carl ROTH), CaCl_2_× 2H_2_O 0.37 g/L (Carl ROTH), MgSO_4_x7H_2_O 0.27 g/L (Sigma-Aldrich).

The cleaned tissues were cut into 1-cm cross sections and embedded into 8% (w/v) low-melting agarose (GERBU, Heidelberg, Germany, #1744-100g)). The agarose was prepared by dissolving the agarose in sterile water and mixing it with an equal volume of double concentrated Roswell Park Memorial Institute medium (RPMI, Gibco, Life Technologies, Paisley, UK, #11544506). The agarose was microwaved until fully dissolved and cooled down to ∼37°C to prevent tissue damage. One-centimetre intestinal sections were embedded in the agarose in a 6-well plate and left to solidify at 4°C for 30 min.

The agarose cores were mounted onto a platform of a Vibratome (Leica VT 1200) and cut into 250 µm-slices at a speed of 0.2 mm/s and an amplitude of 3 mm. The first ten slices were discarded. The PCIS were collected in ice cold KHB buffer and transferred to 24-well plates in 500 µL complete medium (DMEM/Ham’s F12 (Thermo Fisher Scientific, #11320033), 2.5% fetal bovine serum, 10 mg/mL human Insulin (Sigma-Aldrich, #I9278), 1.4 µg/mL human hydrocortisone (Sigma-Aldrich, #H6909), 5 µg/ml human transferrin (Sigma-Aldrich, #T8158), 1 µg/ml rat fibronectin (Sigma-Aldrich, #F0635), 100 U/ml penicillin/100 µg/ml streptomycin (Lonza, Basel, Switzerland), 50 µg/ml gentamicin (Gibco, was excluded for canine PCIS), 1:100 protease inhibitor (Sigma Aldrich, #P8340). The PCIS were incubated at 37 °C in 5% CO_2_ and the medium was exchanged twice with prewarmed complete medium.

To confirm the intestinal section (duodenum, jejunum, ileum), per individual a piece of intestine was collected in ROTI®Histofix 4% Formaldehyde (Carl ROTH, #A146.6) for 24h and used for haematoxylin and eosin staining.

### Testing PCIS viability

PCIS viability was assessed by measuring the adenosine triphosphate (ATP) amount and the total protein content to control for variation in PCIS size as reported previously [37, 53]. For each timepoint, five PCIS were harvested in 1 mL sonication solution (70% Ethanol and 2 mM EDTA), snap frozen and stored at -70°C until further use. Samples were thawed on ice, one Qiagen bead (stainless steel beads 5 mm, #69989) was added per tube and samples were homogenized in a Qiagen Tissuelyser II at 24 Hz for 30 sec. The samples were centrifuged at 13 000 × g for 2 min at 4°C and diluted 1:10 in ATP buffer (100 mM Tris HCl buffer with 2 mM EDTA, pH 7.6). ATP content was measured using the ATP Bioluminescence Assay Kit CLS II (Roche diagnostics, Mannheim, Germany, #11699695001) using an ATP calibration curve according to the manufactureŕs instructions. For the total protein measurement, the remaining pellet was dissolved in 200 µL 5 M NaOH and incubated for 30 min at 37°C. The samples were diluted with 800 µL Milli Q water and the protein content was measured using the BCA kit (Pierce™ BCA Protein Assay Kit, Thermo Fisher Scientific, #23227). For each slice the ratio between ATP content (pmol) and protein content (µg) was calculated.

### Labelling of virus-like particles (VLPs) with Alexa Fluor 488 (AF488)

Virus-like particles have been characterized in previous studies and are listed in **Table 1**. For labelling, Alexa Fluor 488-NHS Ester (Thermo Fisher Scientific; #A20000) was dissolved in DMSO at 10 mg/mL. VLPs were incubated at a 40× molar excess, vortexed and incubated for 1 h at room temperature protected from light. Excess dye was removed by buffer exchange through a 30 kDa-Amicon Filter (Millipore, Burlington MA, USA) at 4200 × g and eluted in PBS. VLP concentration was calculated by Coomassie gel with a BSA standard.

**Table 1:** Virus-like particles.

| **Table 1: Virus-like particles** |  |  |  |  |
| --- | --- | --- | --- | --- |
| **Genotype**<br>(NCBI Accession nr) | **Host** | **Binding to HBGAs** |  | **Ref** |
|  |  | Synthetic glycans | Saliva |  |
| GII.4 Dijon<br>(=US95_96)<br>AF472623.1 | Human | H type 1, A heptasaccharide, Lewis b, Lewis y, Lewis x, SiLe x, Si-LNnT [48] | O Le-, O Le+, A Le-, Ale+, B Le+ [48] | [39] |
| GII.11 (VA34)<br>AY077644 | Pig | None known [54, 55] | None known [54] | [56] |
| GVI.2 (C33)<br>GQ443611.1 | Dog | H type 1 (HSA), A heptasaccharide (HSA), Lewis b (HSA), Sialyl LNF V (HSA) [27] | O Le-, A Le-, O Le+, A Le+, H [27] | [27] |

### Virus histochemistry and immunohistochemistry

The 488-labeled VLPs were used to test for their binding onto formalin-fixed, paraffin-embedded (FFPE) PCIS based on the previously published method [25]. PCIS were kept in 4% Paraformaldehyde for 24 h and were embedded into FFPE blocks and cut into 3-µm slices. To detect VLP attachment the avidin–biotin–peroxidase complex method was used. FFPE slides were deparaffinized 3× 3 min in ROTICLEAR^®^ (Carl ROTH, Karlsruhe, Germany) and subsequently rehydrated through a series of alcohols for 3 min each. Endogenous peroxidase activity was inhibited with 0.5% H_2_O_2_ (in 85% ethanol) for 30 min. Slides were washed three times with PBS and were blocked with 2% bovine serum albumin (BSA) for 30 min. AF488-labeled VLPs were added (500 ng in PBS) and incubated overnight at 4°C. The slides were washed three times with PBS and incubated with a rabbit anti-AF488 antibody (1:100 in PBS with 1% BSA, Thermo Fisher Scientific, #A-11094) for 1 h at RT. The slides were washed three times with PBS and incubated with a biotinylated goat anti-rabbit antibody (1:200 in PBS, Vector Laboratories, Burlingame, CA, USA, #BA-9400) for 45 min at RT. The slides were washed three times with PBS and treated with the avidin–biotin–peroxidase complex (Vectastain ELITE ABC Kit, Vector Laboratories, #PK6100) for 30 min at RT. The slides were washed three times with PBS and the signal was revealed by incubation with 0.05% 3,3′-diaminobenzidine tetrahydrochloride (DAB, Merck KGaA, Darmstadt, Germany) and 0.03% H_2_O_2_ for 5 minutes. The slides were washed three times with tab water followed by 30 seconds nuclear counterstaining with Mayer’s hemalum (Carl ROTH). Slides were washed three times with tab water, dehydrated through a graded alcohol series for 1 min each, followed by 1 min incubation in acetic acid n-butyl ester and xylol and subsequently mounted.

To define the secretor status of each individual through immunohistochemistry, the same protocol was applied as described before but biotin-labelled *Ulex europaeus* agglutinin 1 (UEA1 (1 mg/mL), Sigma-Aldrich, #L8262-2MG,) 1:200 PBS with 1% BSA was incubated over night at 4°C followed by incubation with the biotin amplification complex as described above.

### Inside-out assay

The assay was based on the published protocol [39]; AF488-labelled VLPs (3 µg) were incubated with PCIS or HuTu80 cells (50‘000 seeded/well 24 h prior in 18-well ibiTreat, ibidi, Gräfelfing, Germany, #81816) for 1 h at 4°C. Unbound VLPs were washed away and the PCIS or HuTu80 cells were transferred to 37°C for 2 h, allowing bound VLPs to be internalized. Samples were transferred to ice and from then on treated with pre-cooled solutions on ice. Samples were blocked with DMEM/Ham’s F12 with 2% BSA (blocking buffer) for 25 min and stained for non-internalized VLPs with a rabbit anti-AF488 antibody (1:500, Thermo Fisher Scientific, #A-11094) in blocking buffer for 1 h. Samples were washed three times with PBS and incubated with a secondary goat anti-rabbit AF568 (1:1000, Invitrogen, #A11011) antibody in blocking buffer for 1 h. The plasma membrane and nuclei were stained with AF647-conjugated wheat germ agglutinin (1:500, Thermo Fisher Scientific, #W32466) and Hoechst 33342 (1:5000, Thermo Fisher Scientific, #62249), respectively. Samples were mounted using ProLong™ Diamond Antifade Mountant (Thermo Fisher Scientific, #P36970) and imaged with a Leica SP5 II laser scanning confocal microscope at a 63× magnification. Image settings were consistent for all samples within one experiment. For quantification of VLP entry, at least 100 AF488 events were analysed per individual, VLP, and treatment. The images were analysed using CellProfiler 4.2.6 [57]. VLPs were identified as AF488-positive objects (external and internalized VLPs). The images were masked for AF488 positive objects and overlapping AF568-positive objects (only external VLPs) were identified. The fraction of internalized VLPs was calculated as AF488 objects (all VLPs) - AF488+AF568 (external VLPs) and the percentage of internalized versus all VLPs was calculated.

### Infection of PCIS

After cutting, PCIS were put at 37°C for 1 h to adjust to the temperature. Virus (1×10^7^ genome copies for TGEV and 10^8^ genome copy numbers for HuNoV) was added and incubated for 2 h at 37°C with 5% CO_2_. The inoculum was removed, PCIS were washed with prewarmed complete medium and fresh medium added, supernatant of timepoint 0h was collected. After 24 h, the supernatant was harvested and stored at -70°C. The tissue was put in 4% PFA for immunofluorescence.

### RT-qPCR

For RNA extraction, the QIAamp Viral RNA Mini Kit (Qiagen, Venlo, The Netherlands, #52906) kit was used according to the Manufacturer’s instructions. The samples were prepared for RT-qPCR with the LightCycler® Multiplex RNA Virus Master kit (Roche, #6754155001). RNA (5 µL) was prepared in a 20 µL reaction with 4 µL RT-qPCR Reaction mix, 1 µL RT enzyme solution, 200 nM primers and probe. The following primer and probe pairs were used: For HuNoV GII: QNIF2d: ATGTTCAGRTGGATGAGRTTCTCWGA, COG2R: TCGACGCCATCTTCATTCACA, QNIFS (Probe): FAM-AGCACGTGGGAGGGCGATCG-TAMRA [58]. For TGEV (nucleocapsid) TGEV_F: TGCCATGAACAAACCAAC, TGEV_R: GGCACTTTACCATCGAAT, Probe: HEX-TAGCACCACGACTACCAAGC-BHQ1 [59]. The reaction was performed in LightCycler® 96 System (Roche Diagnostic) under following conditions: For norovirus: reverse transcription at 50°C for 15 min, denaturation at 95°C for 5 min, 40 cycles of amplification with denaturation at 95°C for 15 sec and annealing and extension at 60°C for 35 sec. For TGEV: reverse transcription at 50°C for 30 min, denaturation at 95°C for 15 min, 40 cycles of amplification with denaturation at 94°C for 15 sec and annealing and extension at 60°C for 1 min. Genome copy numbers were calculated per PCIS using RNA standards.

### Immunofluorescent staining

PCIS were fixated in 4% PFA. PCIS were permeabilized in PBS with 0.5% Triton X-100 for 15 min at RT. PCIS were then blocked in PBS with 3% BSA and 0.05% Tween for 60 min at RT. The primary antibodies were eluted 1:100 in PBS with 3% BSA and 0.05% Tween and incubated overnight at 4°C. As primary antibodies we used mouse anti-norovirus capsid protein, mouse anti-coronavirus nucleocapsid protein (Thermo Fisher Scientific, #MA1-82189), mouse anti-double-stranded RNA (J2, Scienion, Berlin, Germany, #10010200). After washing, the secondary antibody (AF488-conjugated goat anti-mouse (Invitrogen, A11001) was added 1:100 in PBS with 0.05% Tween and 3% BSA and incubated for 1 h at RT. Nuclei were stained with Hoechst 33342 (Thermo Fisher Scientific, #62249). PCIS were washed three times with PBS containing 0.05% Tween and mounted in ProLong™ Diamond Antifade Mountant (Thermo Fisher Scientific, #P36970). Images were taken with a Leica SP5 II laser scanning confocal microscope at a 63× magnification.

## Acknowledgments

We thank Paul Becher (Institute of Virology, University of Veterinary Medicine Hannover) and Luis Enjuanes (Centro Nacional de Biotecnología) for providing the TGEV virus and Kerstin Rohn and Melanie Bode for technical advice in the Lab.

This work has been funded by the Deutsche Forschungsgemeinschaft (DFG, German Research Foundation) – 542416050 and the Nationale Forschungsplattform für Zoonosen (Research Network Zoonotic Infectious Diseases) – 01KI2310.

## References

1. Jiang, X., D.Y. Graham, K. Wang, and M.K. Estes, Norwalk virus genome cloning and characterization. Science, 1990. 250(4987): p. 1580–1583.

2. Chhabra, P., M. de Graaf, G.I. Parra, M.C.-W. Chan, K. Green, V. Martella, Q. Wang, P.A. White, K. Katayama, H. Vennema, M.P.G. Koopmans, and J. Vinje, Updated classification of norovirus genogroups and genotypes. J Gen Virol, 2019. 100(10): p. jgv001318.

3. Siebenga, J.J., H. Vennema, D.P. Zheng, J. Vinjé, B.E. Lee, X.L. Pang, E.C.M. Ho, W. Lim, A. Choudekar, S. Broor, T. Halperin, N. Rasool, J. Hewitt, G. Greening, M. Jin, Z.J. Duan, Y. Lucero, M. O’Ryan, M. Hoehne, E. Schreier, R. Ratcliff, P. White, N. Iritani, G. Reuter, and M. Koopmans, Norovirus illness is a global problem: emergence and spread of norovirus GII. 4 variants, 2001–2007. J Infect Dis, 2009. 200(5): p. 802–812.

4. Hoa Tran, T.N., E. Trainor, T. Nakagomi, N.A. Cunliffe, and O. Nakagomi, Molecular epidemiology of noroviruses associated with acute sporadic gastroenteritis in children: Global distribution of genogroups, genotypes and GII.4 variants. Journal of Clinical Virology, 2013. 56(3): p. 269–277.

5. Martino, B.D., F. Di Profio, C. Ceci, E.D. Felice, K.Y. Green, K. Bok, S.D. Grazia, G.M. Giammanco, I. Massirio, E. Lorusso, C. Buonavoglia, F. Marsilio, and V. Martella, Seroprevalence of norovirus genogroup IV antibodies among humans, Italy, 2010-2011. Emerg Infect Dis, 2014. 20(11): p. 1828–1832.

6. Mesquita, J.R., V.P. Costantini, J.L. Cannon, S.C. Lin, M.S.J. Nascimento, and J. Vinjé, Presence of antibodies against genogroup VI Norovirus in humans. Virol J, 2013. 10: p. 176.

7. Villabruna, N., M. Koopmans, and M. De Graaf, Animals as reservoir for human norovirus. Viruses, 2019. 11(5): p. 478.

8. Villabruna, N., R.W. Izquierdo Lara, J. Szarvas, M.P.G. Koopmans, and M.d. Graaf, Phylogenetic investigation of norovirus transmission between humans and animals. Viruses, 2020. 12(11): p. 1287.

9. Charoenkul, K., C. Nasamran, T. Janetanakit, R. Tangwangvivat, N. Bunpapong, S. Boonyapisitsopa, K. Suwannakarn, A. Theamboonler, W. Chuchaona, and Y. Poovorawan, Human Norovirus infection in dogs, Thailand. Emerg Infect Dis, 2020. 26(2): p. 350–353.

10. Summa, M., C.H. von Bonsdorff, and L. Maunula, Pet dogs-A transmission route for human noroviruses? J Clin Virol, 2012. 53(3): p. 244–247.

11. Vinjé, J., M.K. Estes, P. Esteves, K.Y. Green, K. Katayama, N.J. Knowles, Y. L’Homme, V. Martella, H. Vennema, P.A. White, and I.R. Consortium, ICTV Virus Taxonomy Profile: Caliciviridae. J Gen Virol, 2019. 100(11): p. 1469–1470.

12. Summa, M., H. Henttonen, and L. Maunula, Human noroviruses in the faeces of wild birds and rodents-new potential transmission routes. Zoonoses Public Health, 2018. 65(5): p. 512–518.

13. Duarte, M.A., F.J.M. Silva, C.R. Brito, D.S. Teixeira, F.L. Melo, B.M. Ribeiro, T. Nagata, and F.S. Campos, Faecal virome analysis of wild animals from Brazil. Viruses, 2019. 11(9): p. 803.

14. Marionneau, S., N. Ruvoën, B. Le MoullacVaidye, M. Clement, A. CailleauThomas, G. RuizPalacois, P. Huang, X. Jiang, and J. Le Pendu, Norwalk Virus binds to histo-blood group antigens present on gastroduodenal epithelial cells of secretor individuals. Gastroenterology, 2002. 122(7): p. 1967–1977.

15. Hutson, A.M., F. Airaud, J. LePendu, M.K. Estes, and R.L. Atmar, Norwalk virus infection associates with secretor status genotyped from sera. J Med Virol, 2005. 77(1): p. 116–120.

16. Lindesmith, L., C. Moe, S. Marionneau, N. Ruvoen, X.I. Jiang, L. Lindblad, P. Stewart, J. LePendu, and R. Baric, Human susceptibility and resistance to Norwalk virus infection. Nat Med, 2003. 9(5): p. 548–553.

17. Nordgren, J. and L. Svensson, Genetic susceptibility to human norovirus infection: an update. Viruses, 2019. 11(3): p. 226.

18. Hutson, A.M., R.L. Atmar, D.Y. Graham, and M.K. Estes, Norwalk virus infection and disease is associated with ABO histo-blood group type. Emerg Infect Dis, 2002. 185(9): p. 1335–1337.

19. Haga, K., K. Ettayebi, V.R. Tenge, U.C. Karandikar, M.A. Lewis, S.C. Lin, F.H. Neill, B.V. Ayyar, X.L. Zeng, G. Larson, and S. Ramani, Genetic manipulation of human intestinal enteroids demonstrates the necessity of a functional fucosyltransferase 2 gene for secretor-dependent human norovirus infection. mBio, 2020. 11(2): p. 10–1128.

20. Ravn, V. and E. Dabelsteen, Tissue distribution of histo-blood group antigens. APMIS, 2000. 108(1): p. 1–28.

21. Abrantes, J., D. Posada, P. Guillon, P.J. Esteves, and J. Le Pendu, Widespread gene conversion of alpha-2-fucosyltransferase genes in mammals. J Mol Evol, 2009. 69(1): p. 22–31.

22. Le Pendu, J. and N. Ruvoën-Clouet, Fondness for sugars of enteric viruses confronts them with human glycans genetic diversity. Hum Genet, 2020. 139(6): p. 903–910.

23. Thorne, L., A. Nalwoga, A.J. Mentzer, A. de Rougemont, M. Hosmillo, E. Webb, M. Nampiija, A. Muhwezi, T. Carstensen, and D. Gurdasani, The first norovirus longitudinal seroepidemiological study from sub-Saharan Africa reveals high seroprevalence of diverse genotypes associated with host susceptibility factors. Emerg Infect Dis, 2018. 218(5): p. 716– 725.

24. Yamamoto, F., E. Cid, M. Yamamoto, N. Saitou, J. Bertranpetit, and A. Blancher, An integrative evolution theory of histo-blood group ABO and related genes. Sci Rep, 2014. 4: p. 6601.

25. Villabruna, N., C.M. Schapendonk, G.I. Aron, M.P. Koopmans, and M. de Graaf, Human noroviruses attach to intestinal tissues of a broad range of animal species. J Virol, 2020. 95(3): p. e01492–20.

26. Kocher, J.F., L.C. Lindesmith, J. Huynh, J.E. Gates, K. Debbink, A. Beall, M.L. Mallory, E.F. Donaldson, and R.S. Barica, Bat caliciviruses and human noroviruses are antigenically similar and have overlapping histo-blood group antigen binding profiles. mBio, 2018. 9(3): p. e00869–18.

27. Caddy, S., A. Breiman, J. le Pendu, and I. Goodfellow, Genogroup IV and VI canine noroviruses interact with histo-blood group antigens. J Virol, 2014. 88(18): p. 10377–10391.

28. Wang, X., S. Wang, C. Zhang, Y. Zhou, P. Xiong, Q. Liu, and Z. Huang, Development of a surrogate neutralization assay for norovirus vaccine evaluation at the cellular level. Viruses, 2018. 10(1): p. 27.

29. Katayama, K., K. Murakami, T.M. Sharp, S. Guix, T. Oka, R. Takai-Todaka, A. Nakanishi, S.E. Crawford, R.L. Atmar, and M.K. Estes, Plasmid-based human norovirus reverse genetics system produces reporter-tagged progeny virus containing infectious genomic RNA. Proc Natl Acad Sci USA, 2014. 111(38): p. E4043–E4052.

30. Guix, S., M. Asanaka, K. Katayama, S.E. Crawford, F.H. Neill, R.L. Atmar, and M.K. Estes, Norwalk virus RNA is infectious in mammalian cells. J Virol, 2007. 81(22): p. 12238–12248.

31. Brügger, M., C. Machahua, T. Zumkehr, C. Cismaru, D. Jandrasits, B. Trüeb, S. Ezzat, B.I. Oliveira Esteves, P. Dorn, T.M. Marti, G. Zimmer, V. Thiel, M. Funke-Chambour, and M.P. Alves, Aging shapes infection profiles of influenza A virus and SARS-CoV-2 in human precision-cut lung slices. Respir Res, 2025. 26(1): p. 112.

32. Wu, W., W. Zhang, J.L. Booth, and J.P. Metcalf, Influenza A(H1N1)pdm09 virus suppresses RIG-I initiated innate antiviral responses in the human lung. PLoS One, 2012. 7(11): p. e49856.

33. Wolf, S., J. Hewitt, and G.E. Greening, Viral multiplex quantitative PCR assays for tracking sources of fecal contamination. Appl Environ Microbiol, 2010. 76(5): p. 1388–1394.

34. Nguyen, D.T., R.D. de Vries, M. Ludlow, B.G. van den Hoogen, K. Lemon, G. van Amerongen, A.D.M.E. Osterhaus, R.L. de Swart, and W.P. Duprex, Paramyxovirus infections in ex vivo lung slice cultures of different host species. J Virol Methods, 2013. 193(1): p. 159–165.

35. Punyadarsaniya, D., C. Winter, A.K. Mork, M. Amiri, H.Y. Naim, S. Rautenschlein, and G. Herrler, Precision-cut intestinal slices as a culture system to analyze the infection of differentiated intestinal epithelial cells by avian influenza viruses. J Virol Methods, 2015. 212: p. 71–5.

36. Krimmling, T., A. Beineke, and C. Schwegmann-Weßels, Infection of porcine precision cut intestinal slices by transmissible gastroenteritis coronavirus demonstrates the importance of the spike protein for enterotropism of different virus strains. Vet Microbiol, 2017. 205: p. 1–5.

37. de Graaf, I.A.M., P. Olinga, M.H. de Jager, M.T. Merema, R. de Kanter, E.G. van de Kerkhof, and G.M.M. Groothuis, Preparation and incubation of precision-cut liver and intestinal slices for application in drug metabolism and toxicity studies. Nat Protoc, 2010. 5(9): p. 1540–1551.

38. Kerkhof, E.G.v.d., I.A.M.d. Graaf, A.-L.B. Ungell, and G.M.M. Groothuis, Induction of metabolism and transport in human intestine: validation of precision-cut slices as a tool to study induction of drug metabolism in human intestine in vitro Drug Metab Dispos, 2008. 36(3): p. 604–613.

39. Palm, E., K. Danskog, S. Nord, M. Becker, H. Tanner, L. Sandblad, D. Öhlund, A. Lenman, and N. Arnberg, Bile acids accumulate norovirus-like particles and enhance binding to and entry into human enteric epithelial cells. bioRxiv, 2023: p. 2023.04.28.538707.

40. Almand, E.A., M.D. Moore, J. Outlaw, and L.A. Jaykus, Human norovirus binding to select bacteria representative of the human gut microbiota. PLoS One, 2017. 12(3): p. e0173124.

41. Li, D., A. Breiman, J. le Pendu, and M. Uyttendaele, Binding to histo-blood group antigen-expressing bacteria protects human norovirus from acute heat stress. Front Microbiol, 2015. 6: p. 659.

42. Gao, X., M.A. Esseili, Z.Y. Lu, L.J. Saif, and Q.H. Wang, Recognition of histo-blood group antigen-like carbohydrates in lettuce by human GII.4 norovirus. Appl Environ Microbiol, 2016. 82(10): p. 2966–2974.

43. Wang, M., S.F. Rong, P. Tian, Y. Zhou, S.M. Guan, Q.Q. Li, and D.P. Wang, Bacterial surface-displayed GII. 4 human norovirus capsid proteins bound to HBGA-like molecules in romaine lettuce. Front Microbiol, 2017. 8: p. 251.

44. Tian, P., A.L. Engelbrektson, X. Jiang, W. Zhong, and R.E. Mandrell, Norovirus recognizes histo-blood group antigens on gastrointestinal cells of clams, mussels, and oysters: a possible mechanism of bioaccumulation. J Food Prot, 2007. 70(9): p. 2140–7.

45. Maalouf, H., J. Schaeffer, S. Parnaudeau, J.L. Pendu, R.L. Atmar, S.E. Crawford, and F.S.L. Guyader, Strain-dependent norovirus bioaccumulation in oysters. Appl Environ Microbiol, 2011. 77(10): p. 3189–3196.

46. White, L.J., J.M. Ball, M.E. Hardy, T.N. Tanaka, N. Kitamoto, and M.K. Estes, Attachment and entry of recombinant Norwalk virus capsids to cultured human and animal cell lines. J Virol, 1996. 70(10): p. 6589–6597.

47. Oka, T., G.T. Stoltzfus, C. Zhu, K. Jung, Q. Wang, and L.J. Saif, Attempts to grow human noroviruses, a sapovirus, and a bovine norovirus in vitro. PLoS One, 2018. 13(2): p. e0178157.

48. Rougemont, A.d., N. Ruvoen-Clouet, B. Simon, M. Estienney, C. Elie-Caille, S. Aho, P. Pothier, J.L. Pendu, W. Boireau, and G. Belliot, Qualitative and quantitative analysis of the binding of GII. 4 norovirus variants onto human blood group antigens. J Virol, 2011. 85(9): p. 4057–4070.

49. Ettayebi, K., S.E. Crawford, K. Murakami, J.R. Broughman, U. Karandikar, V.R. Tenge, F.H. Neill, S.E. Blutt, X.L. Zeng, L. Qu, B. Kou, A.R. Opekun, D. Burrin, D.Y. Graham, S. Ramani, R.L. Atmar, and M.K. Estes, Replication of human noroviruses in stem cell–derived human enteroids. Science, 2016. 353(6306): p. 1387–1392.

50. Ayyar, B.V., K. Ettayebi, W. Salmen, U.C. Karandikar, F.H. Neill, V.R. Tenge, S.E. Crawford, E. Bieberich, B.V.V. Prasad, R.L. Atmar, and M.K. Estes, CLIC and membrane wound repair pathways enable pandemic norovirus entry and infection. Nat Commun, 2023. 14(1): p. 1148.

51. Ettayebi, K., V.R. Tenge, N.W. Cortes-Penfield, S.E. Crawford, F.H. Neill, X.-L. Zeng, X. Yu, B.V. Ayyar, D. Burrin, S. Ramani, R.L. Atmar, M.K. Estes, and S.M.M. Esstman, New insights and enhanced human norovirus cultivation in human intestinal enteroids. mSphere, 2021. 6(1): p. e01136–20.

52. Pohl, C., G. Szczepankiewicz, and U.G. Liebert, Analysis and optimization of a Caco-2 cell culture model for infection with human norovirus. Archives of Virology, 2022.

53. Bigaeva, E., J.J.M. Bomers, C. Biel, H.A.M. Mutsaers, I.A.M. de Graaf, M. Boersema, and P. Olinga, Growth factors of stem cell niche extend the life-span of precision-cut intestinal slices in culture: A proof-of-concept study. Toxicol in Vitro, 2019. 59: p. 312–321.

54. Yang, Y., M. Xia, L. Wang, S. Arumugam, Y. Wang, X. Ou, C. Wang, X. Jiang, M. Tan, C. Y., and L. X., Structural basis of host ligand specificity change of GII porcine noroviruses from their closely related GII human noroviruses. Emerg Microbes Infect, 2019. 8(1): p. 1642–1657.

55. Farkas, T., S. Nakajima, M. Sugieda, X. Deng, W. Zhong, and X. Jiang, Seroprevalence of noroviruses in swine. J Clin Microbiol, 2005. 43(2): p. 657–661.

56. Almanza, H., C. Cubillos, I. Angulo, F. Mateos, J.R. Castón, W.H.M. Van Der Poel, J. Vinje, J. Bárcena, and I. Mena, Self-assembly of the recombinant capsid protein of a swine norovirus into virus-like particles and evaluation of monoclonal antibodies cross-reactive with a human strain from genogroup II. J Clin Microbiol, 2008. 46(12): p. 3971–3979.

57. Stirling, D.R., M.J. Swain-Bowden, A.M. Lucas, A.E. Carpenter, B.A. Cimini, and A. Goodman, CellProfiler 4: improvements in speed, utility and usability. BMC Bioinformatics, 2021. 22(1): p. 433.

58. Loisy, F., R.L. Atmar, P. Guillon, P. Le Cann, M. Pommepuy, and F.S. Le Guyader, Real-time RT-PCR for norovirus screening in shellfish. J Virol Methods, 2005. 123(1): p. 1–7.

59. Huang, X., J. Chen, G. Yao, Q. Guo, J. Wang, and G. Liu, A TaqMan-probe-based multiplex real-time RT-qPCR for simultaneous detection of porcine enteric coronaviruses. Appl Microbiol Biotechnol, 2019. 103(12): p. 4943–4952.

